# A Chemically-triggered Transition from Conflict to Cooperation in Burying Beetles

**DOI:** 10.1101/389163

**Authors:** Bo-Fei Chen, Mark Liu, Dustin R. Rubenstein, Syuan-Jyun Sun, Jian-Nan Liu, Yu-Heng Lin, Sheng-Feng Shen

## Abstract

Although interspecific competition has long been recognized as a major driver of trait divergence and adaptive evolution^1–3^, relatively little effort has focused on how it influences the evolution of intraspecific cooperation^4–6^. Here we identify the mechanism by which the perceived pressure of interspecific competition influences the transition from intraspecific conflict to cooperation in a facultative cooperatively breeding species, the Asian burying beetle *Nicrophorus nepalensis*. In their natural environment in central Taiwan, *N. nepalensis* are typically aggressive to conspecifics and only cooperate with others of their own species at critical carcass resources in the presence of blowflies, their primary competitors^7^. We demonstrate that beetles form larger groups and are more cooperative in carcass preparation in warmer environments where the pressure of interspecific competition with blowflies is highest^8^. To test the hypothesis that the presence of blowflies promotes beetle cooperation and to identify the mechanism by which this occurs, we manipulated blowfly larvae on carcasses in the lab. We not only found that beetles are more cooperative at carcasses when blowfly maggots have begun to digest the tissue, but that this social cooperation appears to be triggered by a single chemical cue— dimethyl disulfide (DMDS)—emitted from carcasses consumed by blowflies but not from control carcasses lacking blowflies. Our results provide experimental evidence that interspecific competition promotes the transition from intraspecific conflict to cooperation in *N. nepalensis* via a surprisingly simple social chemical cue that is a reliable indicator of interspecific competition. This finding helps bridge the gap between the proximate and ultimate factors regulating the transition between cooperation and conflict and moves toward a more comprehensive understanding of the evolution of mechanisms governing intraspecific variation in social behaviour.

---

Burying beetles (*Nicrophorus spp.*) use small vertebrate carcasses as their sole resource for reproduction and often face intense intra- and interspecific competition for access to these precious but limiting resources^9–11^. Previous work has suggested that the key benefit of cooperation in the Asian burying beetle *N. nepalensis* is that cooperative carcass preparation—including carcass cleaning, shaping, and burial, as well as the elimination of competing species^9–12^—enables beetles to outcompete their primary competitor, blowflies (family Calliphoridae), particularly in warmer environments where blowflies are most abundant. By experimentally manipulating burying beetle group size along an elevational gradient, we showed that in cooler environments where the pressure of interspecific competition is low, beetles in large groups are more aggressive toward same-sex conspecifics and often engage in intense and even lethal fights that result in a single individual monopolizing the carcass and having a higher probability of breeding successfully than those in large groups^8^. In contrast, in warmer environments where blowflies are more common, burying beetles cooperate with conspecifics to more quickly bury carcasses and escape blowfly competition^7^, ultimately gaining greater reproductive success^8^. Although the presence of blowflies at carcasses appears to facilitate a shift from competitive to cooperative behaviour in *N. nepalensis*, it remains unclear what drives this transition in beetle social behaviour and how individuals know to reduce conflict and tolerate conspecifics.

To determine how ecology influences inter- and intraspecific social interactions in natural burying beetle populations, we first quantified beetle social behaviour and dynamics by video recording their breeding behaviours at 25 sites along two elevational gradients in eastern and western Taiwan, each spanning more than 1000 m in elevation. We calculated the time that beetles spend on cooperative carcass preparation (hereafter cooperative investment) both in terms of total investment (i.e. the cumulative time of the social group) and on a per capita basis for large (groups larger than the median size) and small groups (groups smaller than the median size), as well as in cool (<14.5°C) and warm environments (>14.5°C). We found that group size peaked at moderate temperatures (χ^2^_1_ = 5.52, *P* = 0.019, n = 245; Fig. 1a) and that per capita cooperative investment along the temperature gradient varied with group size (group size × temperature interaction, χ^2^_1_ = 11.20, *P* = 0.001, *n =* 89). Specifically, per capita cooperative investment increased with daily minimum temperature in large groups (χ^2^_1_ = 5.39, *P* = 0.02, *n* = 33), but not in small groups (χ^2^_1_ = 0.05, *P* = 0.83, *n =* 56; Fig 1b). Similarly, total cooperative investment increased with daily minimum temperature in large groups (χ^2^_1_ = 4.88, *P* = 0.03, *n* = 33), but not in small groups (χ^2^_1_ = 0.24, *P* = 0.60, *n =* 56; Fig. 1c). In contrast, per capita social conflict along the temperature gradient, measured as the number of intraspecific conflict events for each individual, varied with group size (mean group size × temperature interaction, χ^2^_1_ = 6.64, *P* < 0.01, *n =* 82, Fig. 2b), such that conflict increased with group size in cool environments (χ^2^_1_ = 11.24, *P* < 0.001, *n =* 40; Fig. 1d), but not in warm environments (χ^2^_1_ = 1.59, *P* = 0.2, *n =* 42; Fig. 1d).

**Figure 1.**
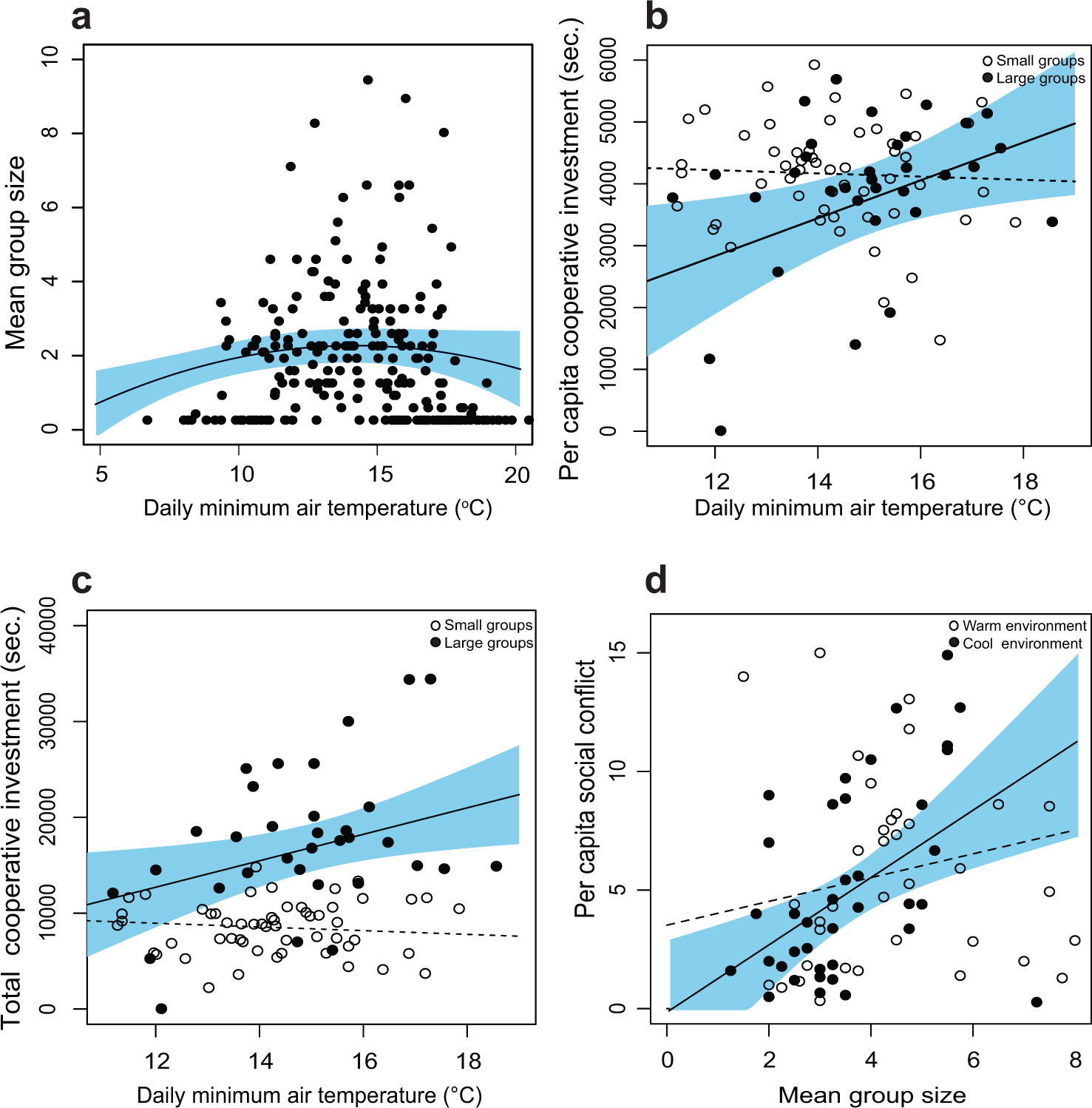
Changes in *N. nepalensis* group size and social behaviours during carcass preparation along a temperature gradient. The relationship between daily minimum air temperature and **(a)** mean group size, **(b)** per capita cooperative investment, **(c)** total cooperative investment in large (closed circles) and small groups (open circles). Group size peaked at moderate temperatures, whereas per capita and total cooperative investment increased with daily minimum temperature in large but not small groups. Solid lines denote predicted relationships from GLMs, whereas dashed lines denote non-significant relationships. **(d)** Per capita social conflict increased with group size in cool environments (closed circles), but not in warm environments (open circles). Lines represent least-squared means (solid lines denote significant relationships and dotted lines non-significant relationships), and blue shaded areas represent 95% confidence intervals expected from GLMMs.

**Figure 2.**
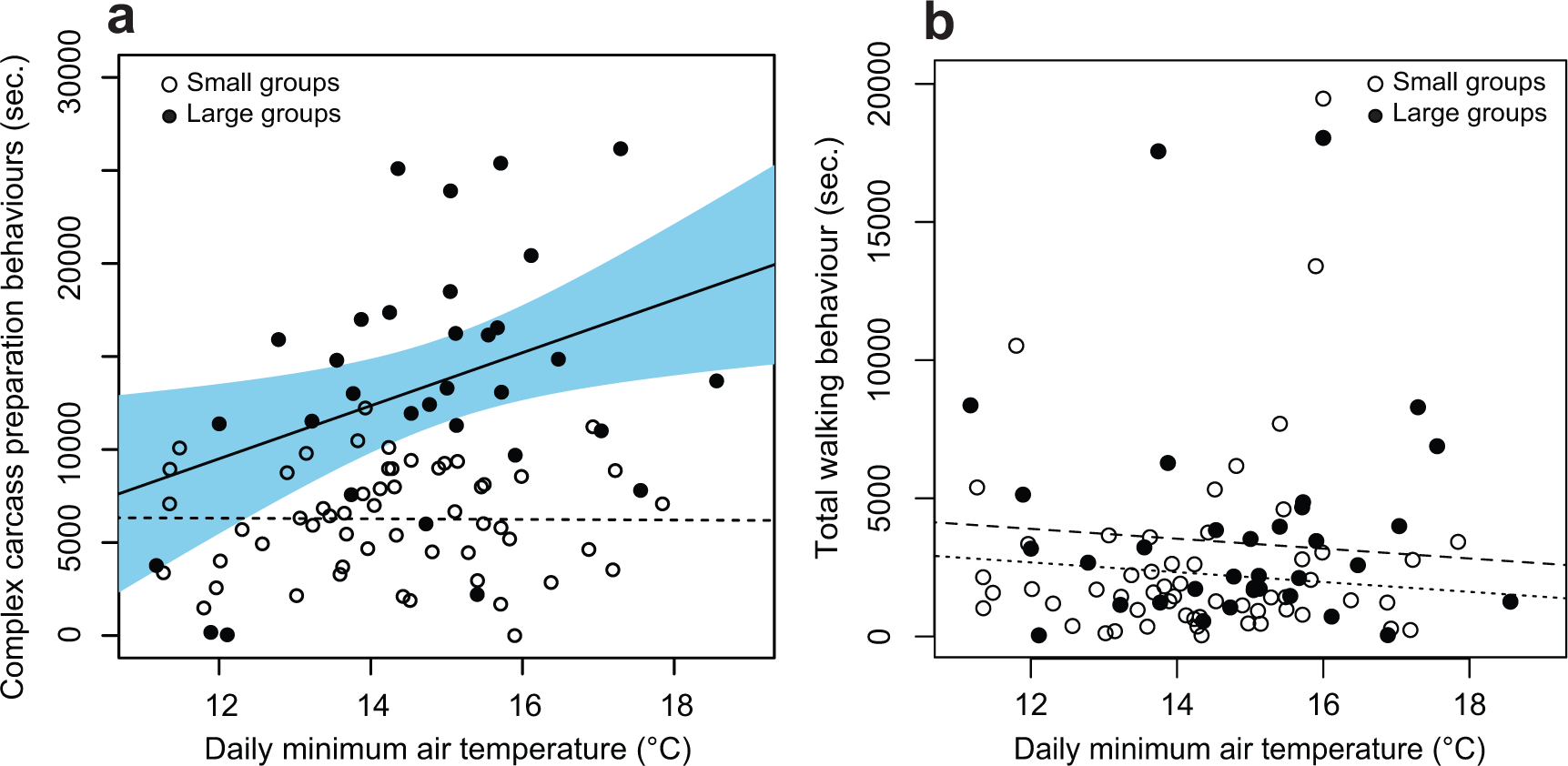
Complex carcass preparation and simple walking behaviours during cooperative carcass preparation along the temperature gradient. The time that beetles spent on **(a)** complex carcass preparation behaviours and **(b)** walking on the carcass in relation to daily minimum air temperature in large and small groups. Compared to small groups (open circles), large groups (closed circles) spent more time on complex carcass preparation but not on walking as daily minimum air temperature increased, suggesting that the increase in total cooperative investment in warmer environments was not simply the result of increased activity at warmer temperatures. Lines represent least-squared means (solid lines denote significant relationships and dotted lines non-significant relationships), and the blue shaded area represent 95% confidence intervals expected from GLMMs.

To confirm that these patterns of social conflict and cooperation were the result of changes in social behaviour and not simply changes in activity associated with differences in ambient temperature, we further separated cooperative investment into (1) time spent simply walking on the carcass and (2) more complex carcass-preparation behaviours, which are presumably more costly— including maggot and rotten tissue removal, as well as carcass dragging, depilation, and burial. We found that time spent on more complex carcass preparation behaviours increased with increasing daily minimum temperature in large groups (χ^2^_1_ = 5.39, *P* = 0.02, n = 33; Fig. 2a), but not in small groups (χ^2^_1_ = 0.17, *P* = 0.68, n = 56; Fig. 2a). However, there was no significant relationship between walking time and daily minimum temperature in large (χ^2^_1_ = 0.24, *P*= 0.60, n = 33; Fig. 2b) or small groups (χ^2^_1_ = 0.79, *P* = 0.37, n = 56; Fig. 2b), suggesting that the increase in total cooperative investment in warmer environments was not simply the result of increased activity at warmer temperatures.

Our field results demonstrate that *N. nepalensis* exhibits remarkably flexible social behaviours along elevational and thermal gradients: beetles are normally asocial and aggressive towards conspecifics in cooler environments but become social and cooperate with conspecifics in warmer environments where the competition for critical resources with other species is intense^7^. However, to demonstrate experimentally that blowfly competition for carcasses drives the transition from intraspecific competition to intraspecific cooperation, we performed a series of laboratory experiments to directly manipulate the presence or absence of blowflies at carcasses.

Our first experiment introduced blowfly competition to burying beetles by exposing carcasses to adult blowflies in an incubator at 26°C for two days, conditions that match those in the field and are optimal for blowflies to lay eggs and for their maggots to partially consume the carcass. We then allowed six beetles (three males and three females) to breed on the carcass. We found that more beetles cooperated (t = 5.26, *P* < 0.001; Fig. 3a), and that each individual beetle spent significantly more time cooperating, in the blowfly treatment than in the control treatment containing carcasses but no blowflies (t = 3.27, *P* = 0.002; Fig. 3b). As a consequence, the total cooperative investment was higher in the blowfly treatment than in the control treatment (t = 5.37, *P* < 0.001; Fig. 3c). Although there was no difference in per capita social conflict between the blowfly and control treatments (t = −0.33, *P* = 0.75; Fig. 3d), after controlling for total investment time by dividing per capita social conflict by the total cooperative investment, the adjusted per capita number of social conflicts per unit time was significantly lower in the blowfly treatment than in the control treatment (t = - 2.58, *P* = 0.013; Fig. S1). Thus, social conflict in burying beetles was lower and cooperation higher when blowflies were present on carcasses.

**Figure 3.**
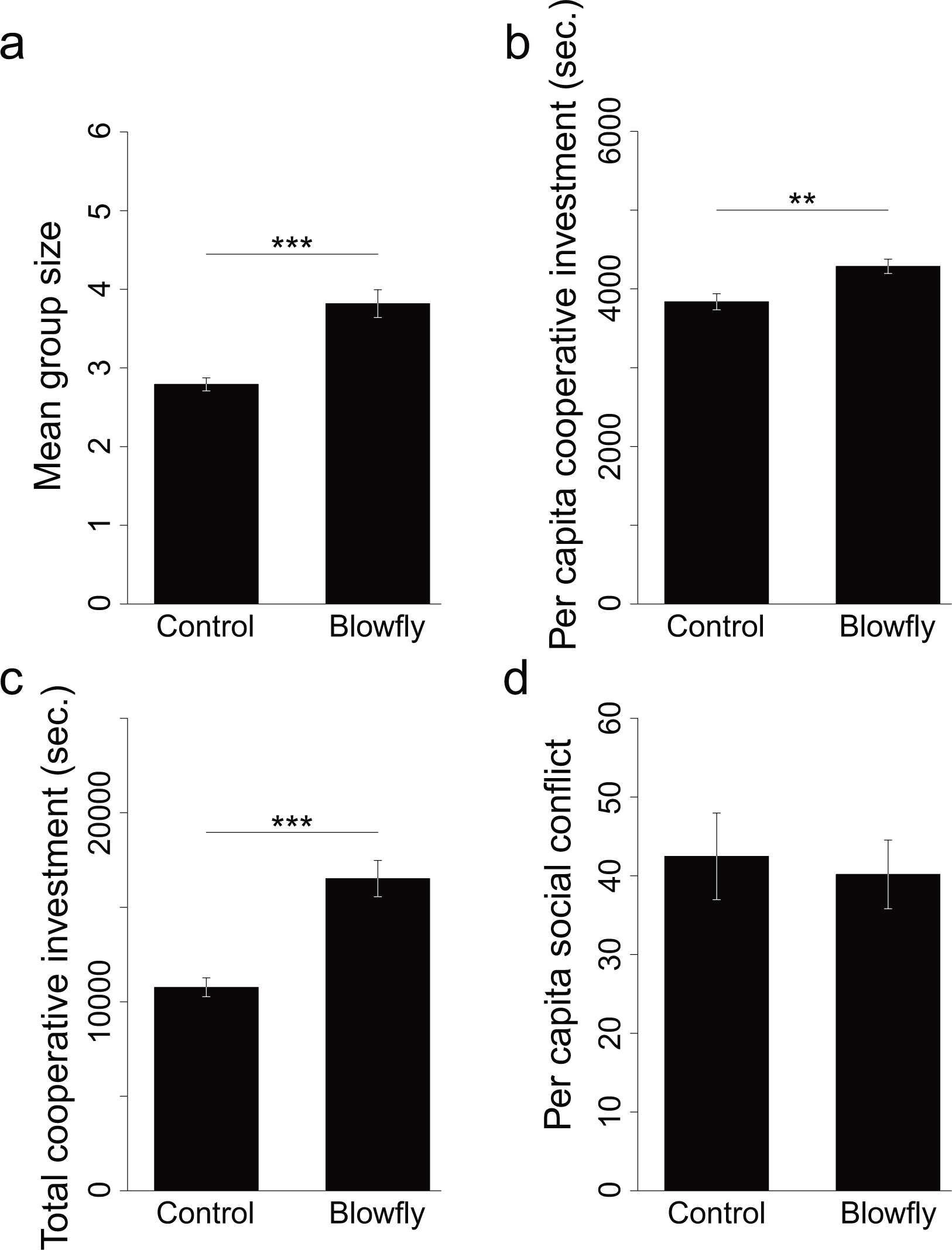
*N. nepalensis* social behaviours in control and blowfly treatments. **(a)** Mean group size, **(b)** per capita cooperative investment, **(c)** total cooperative investment, and **(d)** per capita social conflict of burying beetles on carcasses. Beetles formed larger groups and had greater per capital and total cooperative investment in carcass preparation in the presence of blowflies than in control treatments where blowflies were absent. ** *P* ≤ 0.01; *** *P* ≤ 0.001.

What is the mechanism driving the transition from intraspecific competition to intraspecific cooperation? Since blowfly species are diurnal but *N. nepalensis* is nocturnal, it is unlikely that the physical presence of blowflies influences *N. nepalensis* behaviour. Previous studies have demonstrated that sulfur-containing volatile organic compounds (S-VOCs) emitted from decomposing carcasses attract burying beetles to this key resource^13,14^. Because GC-MS analysis showed that dimethyl disulfide (DMDS) appeared earlier and was more abundant in the blowfly treatment than in the control (Fig. 4a), we hypothesized that DMDS is the key infochemical^15^—indicating not only the presence of a decaying carcass but also the degree of interspecific competition at that carcass—that mediates the transition between cooperative and competitive strategies in *N. nepalensis*.

**Figure 4.**
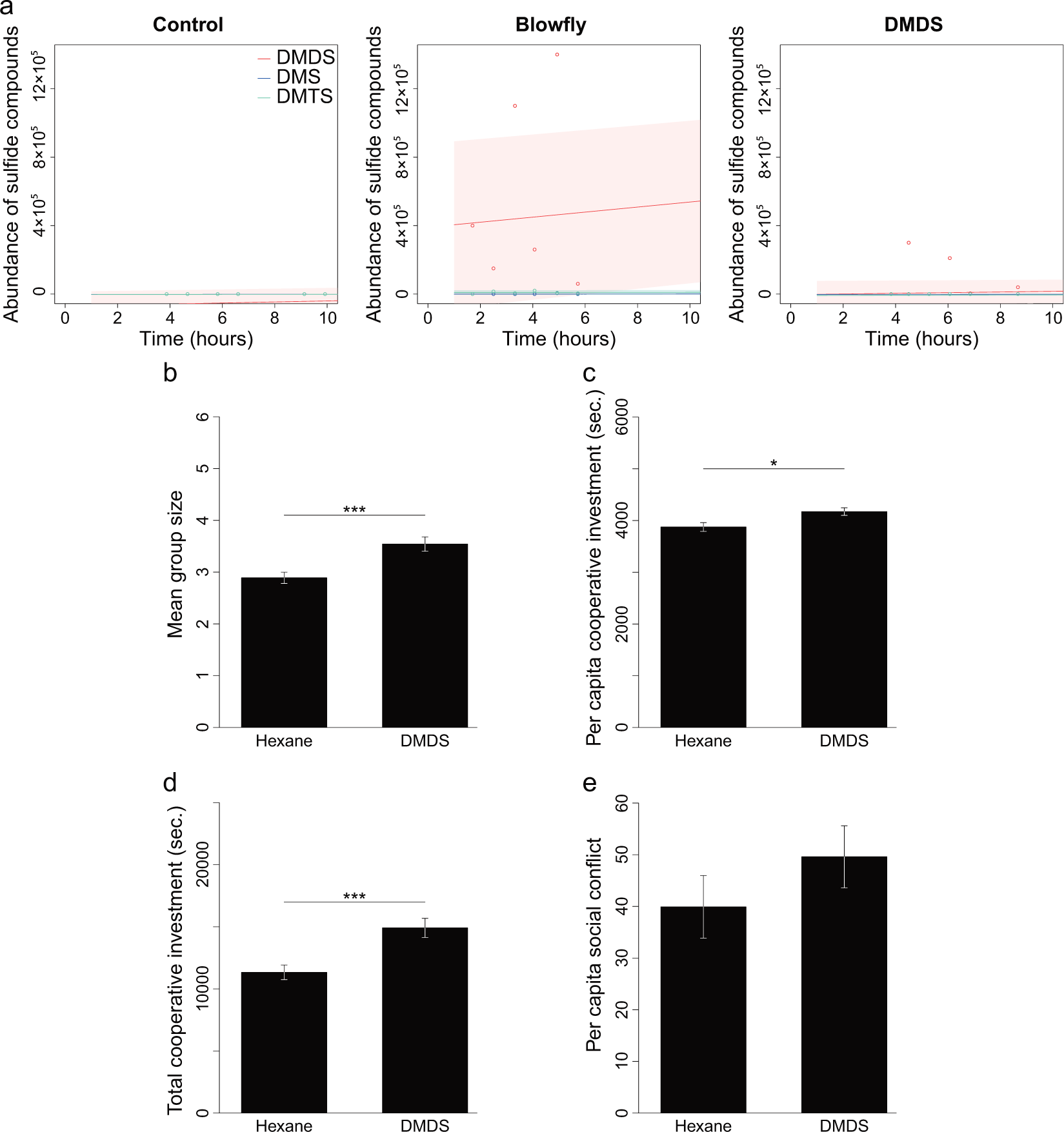
Results of gas chromatography-mass spectrometry (GC-MS) analyses and *N. nepalensis* social behaviours in hexane and DMDS treatments. **(a)** GC-MS analyses showed an abundance of sulfide compounds, including dimethyl sulfide (DMS), dimethyl disulfide (DMDS) and dimethyl trisulfide (DMTS) in control, blowfly, and DMDS treatments during the first 10hrs. DMDS was the major sulfide compound emitted by maggot-digested carcasses. Shaded areas represent 95% confidence intervals expected from GLMMs. **(b)** Mean group sizes, **(c)** per capita cooperative investment, **(d)** total cooperative investment, and **(e)** per capita social conflict of burying beetles on carcasses in DMDS and hexane control treatments. Beetles formed larger groups and had greater per capital and total cooperative investment on carcasses in the DMDS treatment compared to the hexane control treatment. * *P* ≤ 0.05; *** *P* ≤ 0.001.

To experimentally test this hypothesis, we injected DMDS into the body cavity of mouse carcasses. We found that more individuals cooperated (t = −3.76, *P* < 0.001; Fig. 4b), and that each individual spent more time cooperating, in DMDS treated carcasses relative to controls (t = −2.55, *P* = 0.014; Fig. 4c). Thus, there was a higher total cooperative investment in the DMDS treatment than in the control (t = −3.8, *P* < 0.001; Fig. 4d). These results were similar to those observed in the blowfly treatment from the initial experiment. The only difference between the DMDS and blowfly treatments was that there was marginally more social conflict in the DMDS treatment than in the hexane control (t = −1.97, *P* = 0.054; Fig. 4e), whereas this trend—while in the same direction—was not significant in the blowfly treatment (t = −0.33, *P* = 0.75), presumably because there were no real competitors that beetles need to remove in the DMDS treatment.

Our study shows that burying beetles transition from competitive to more cooperative interactions as the pressure of interspecific competition increases. Accumulating empirical evidence from other animals suggests that social conflict in cooperative societies is often lower in adverse environments with strong interspecific competition^5^. This pattern of reduced social conflict under strong interspecific competition has largely been explained by the fact that the cost of engaging in competitive interactions increases under adverse conditions^16^. Yet, there is little empirical evidence demonstrating that social animals increase their investment in cooperation under the threat of interspecific competition. One exception comes from cooperatively breeding superb fairy-wrens (*Malurus cyaneus*) who cooperate more in nest defence when exposed to a greater threat of interspecific brood parasitism^17^. However, it remains unclear how intraspecific conflict in fairy-wrens is influenced by the threat of interspecific competition. Our study helps fill this knowledge gap by showing that cooperative carcass preparation to reduce blowfly competition in warmer environments is critical for predicting both the cooperative and competitive interactions among individuals of the same species.

Furthermore, we show that the conditional cooperative and competitive strategies^18,19^ used by *N. nepalensis* to maximize their utility of carcasses are mediated by a surprisingly simple chemical mechanism. DMDS is produced during the decomposition process and acts as an indicator of the level of competition from blowflies. Although interspecific competition has long been recognized as a major ecological force that drives adaptive evolution^1–3^, relatively little effort has focused on how it influences intraspecific cooperation^4–6^. Our discovery of a novel social chemical cue provides unambiguous evidence that interspecific competition has shaped social evolution in *N. nepalensis*. DMDS acts as a kairomone because it is produced by heterospecifics (i.e. blowfly digestion), but benefits the receiver^15^, and not as a pheromone produced by conspecifics^20,21^. Pheromones are often used for kin discrimination, and studying the olfactory sensory system and its genes have greatly advanced our understandings of the role that chemically-driven kin recognition has played in social evolution, especially in ants^22,23^. Here we demonstrate that interspecific chemical communication is also important to insect social evolution. Ultimately, by showing that chemically-mediated interspecific competition is a key driver of intraspecific cooperation and of social evolution more generally, our work demonstrates the value of integrating ultimate and proximate levels to study the evolution of cooperation^24^.

## Methods

### Field Study

The field study was conducted in Taiwan from 2012 to 2015 along elevational gradients composed primarily of uncultivated forest in Nantou county (spanning 1688 m to 2650 m above sea level) and Hualien county (spanning 1193 m to 2720 m above sea level). A variety of social behaviours, including per capita social conflict and investment in cooperative carcass preparation (i.e. cooperative investment), were scored on the first night of video observation (from 19:00 to 05:00) using the Observer^®^ XT 14 (Noldus). To quantify total cooperative investment, we estimated the cumulative time that each beetle spent depilating rat hair, cleaning rat carcasses by removing maggots, or dragging carcasses during carcass burial and preparation. We measured per capita cooperative investment as the total cooperative investment divided by the mean group size, defined as the maximum number of beetles that stayed on or under carcasses averaged over three time periods (22:30 to 22:40; 01:30 to 01:40 and 04:30 to 04:40) during each video observation. Investment in cooperation was quantified as the duration of cumulative time sampled for a 10 mins observation period in each hour (i.e. 100 mins for each breeding experiment). In total, there were 89 breeding experiments (resulting in 8900 mins of video recordings) from which we were able to quantify total cooperative investment. Aggressive interactions were defined as social conflict if a beetle attacked, wrestled, chased, or escaped from another same-sexed individual (see below for definitions of each behaviour). We measured total social conflict as the total number of aggressive interactions over the 240 mins observation period. We measured per capita social conflict as the total number of aggressive interactions divided by the mean group size for each observation period. Conflict was quantified as the total number of aggressive interactions sampled for two 120 mins observation periods (from 19:30 to 21:30 and from 23:30 to 1:30). In total, there were 82 breeding experiments (resulting in 19,680 mins of video recordings) from which we were able to quantify conflict behaviour. We determined the mean group size on the first night of each beetle’s arrival in 245 breeding experiments (resulting in 7350 mins of video recordings).

### Collection and maintenance

Lab experiments were carried out using *N. nepalensis* individuals from laboratory-reared strains that originated from Meifeng, Nantou County, Taiwan (24°5’ N, 121°10’). Burying beetles were collected using hanging pitfall traps baited with 100 g rotten chicken breasts. Collected beetles were randomly paired and supplied with frozen and re-thawed 75 ± 5g dead rats (*Rattus norvegicus*) in 23 × 15.5 × 16 cm plastic boxes filled with 10 cm moist peat for reproduction. The emerged beetles were housed individually in 7.3 × 7.3 × 3.5 cm plastic boxes filled with 2 cm moist peat and fed with dead superworms (*Zophobas morio*) once a week. All individuals were kept in environmental chambers at 13.2 ~ 19.7 °C (to resemble the natural daily temperature fluctuation in their natural habitat) on a 14 L:10 D photoperiod. Experimental beetles were between 40 and 80 days of age, which is their optimal age for reproduction (individuals can live for over three months in the laboratory).

### Experimental design and procedure

For each experimental replicate, three unrelated males and three unrelated females were randomly chosen from different families to avoid relatedness affecting their behaviours. Each individual was weighed to the nearest 0.1 mg and marked with a Uni POSCA paint marker on the elytra and coated with Scorch^®^ Super GlueGel for individual identification in videos. The marking and weighing of beetles was done 2 hrs prior to beginning an experiment to ensure that all beetles would return to normal activity levels. All six marked beetles were placed into the experimental boxes in random order at the beginning of each experiment. Experimental boxes consisted of a smaller plastic container (23 × 15.5 × 13.5 cm filled with 13.5 cm moist peat) located inside a larger plastic container (45 × 34.5 × 25 cm filled with 13.5 cm moist peat). A 4 cm high iron net with 2 cm^2^ mesh was placed around the small container to prevent beetles moving carcasses outside the field of view of the digital cameras, but beetles could still move freely between the inner and outer areas. A digital camera was fitted on the top of a 25 × 20 × 55 cm acrylic box, which was fixed on the cap of the large container. To equalize the temperature of the experimental apparatus, boxes were filled with moist peat and put into the environmental chambers one day before the experiments began.

The blowfly treatment was conducted by exposing a 75 ± 5 g rat thawed carcass to blowflies, oriental latrine flies (*Chrysomya megacephala*), in 32 × 32 × 32 fly cages for 50 hrs before the start of each experiment. Fly cages contained oriental latrine flies that had emerged from 10 g pupa and been kept in environmental chambers at 26°C on a 14 L:10 D photoperiod. Except for maggot-digested carcasses, all other carcasses in the same weight range were thawed at 4°C for 24 hrs before experiments began. Carcasses used in all treatments were moved into the environmental chambers 8 hrs prior to the start of experiments to equalize their temperatures. The hexane control and DMDS treatment used thawed-only carcasses injected with 2 ml hexane or 0.01 M DMDS solution, respectively, into abdominal cavities through the anus using 3 ml Terumo^®^ Syringes and needles 1 hr prior to the start of the experiment. The thawed-only carcasses served as controls. The carcasses in controls and all treatments were moved into the experimental boxes and put on the surface of peat in smaller containers 1 hr before experiments began. Behavioural videos were recorded either from 7 PM until the day and time at which a carcass was completely buried into peat or for 72 hrs if the beetles did not completely bury the carcass (under natural conditions, a carcass would be completely consumed by blowflies if beetles did not completely bury it within 72 hrs). In total, 1020 hrs of videos were analyzed from 23 blowfly control replicates, 23 blowfly treatment replicates, 32 hexane control replicates, and 24 DMDS replicates. Social conflict and cooperative investment behaviours were recorded in the first 10 hrs (7 PM to 5 AM) of each experimental treatment using The Observer^®^ XT 14 (Noldus).

### Gas chromatography-mass spectrometry (GC-MS) analysis

The composition of volatile organic compounds (VOCs) emitted by carcasses was determined from two control and two blowfly treated carcasses prepared using the same procedure described previously. The prepared carcasses were put on the peat surface in glass vacuum desiccators (15 cm diameter × 22 cm tall) filled with 5 cm of moist peat. The stopcock and ground-glass rim of the desiccator lid were greased with a thin layer of petroleum jelly to prevent the leakage of emitted VOCs, as well as contamination from the atmosphere. The VOCs were sampled using solid-phase micro-extraction (SPME)^25^. The SPME holder with CAR/PDMS fiber (Supelco, previously desorbed for 5 mins in GC injection port heated to 200°C) was inserted through the hole of the stopcock into the atmosphere surrounding the rat carcass. Immediately after exposing the fiber for 15 mins, the sample was GC-MS-analyzed using a 6890N Network Gas Chromatograph (Agilent Technologies) equipped with a HP-5ms column (Agilent J&W) and a 5975 Mass Selective Detector (Agilent Technologies). The GC oven was operated at an initial temperature 40°C for 1 min and then ramped up at a rate 10°C per min to 250°C (with a 10 mins hold). The temperatures of the GC inlet and detector were set to 200°C and 260°C, respectively. The SPME samples were GC analysed split-less. Helium (1 ml per min) was used as a carrier gas. Since the GC-MS results showed DMDS was the major VOC emitted by the blowfly-treated carcasses, DMDS was injected into carcasses in the further experiments. Two DMDS-injected carcasses (also prepared using the same procedure described previously) were used in GC-MS analyses (following the procedure described above) to determine the composition of the VOCs they emitted.

### Statistical analyses

Multivariate analyses were performed using generalized linear models (GLMs) to determine statistical significance for differences between controls and blowfly treatments or hexane controls and DMDS treatments in mean group size, total and per capita cooperative investment, and total and per capita social conflict. All statistical analyses were performed in R using the packages stats, lme4, car, multcomp (http://cran.r-project.org/), and glmmADMB (http://glmmadmb.r-forge.r-project.org/).

## Author contributions

S.-F.S. conceived the idea for the study. B.-F.C., M.L., D.R.R. and S.-F.S. design the experiments. B.-F.C., M.L., S.-J.S. performed field experiment. B.-F.C. and J.-N.L. performed lab behavior experiments. B.-F.C. and Y.-H.L. did the GC-MS analysis. B.-F.C., M.L., D.R.R. and S.-F.S. analyzed the data and wrote the paper.

## Acknowledgements

S.-F.S was supported by Career Development Award and Investigator Award, Academia Sinica and Ministry of Science and Technology of Taiwan (100-2621-B-001-001, 103-2621-B-001-003-MY3). D.R.R. was supported by the US National Science Foundation (IOS-1257530 and IOS-1656098).

---

**Figure S1:**
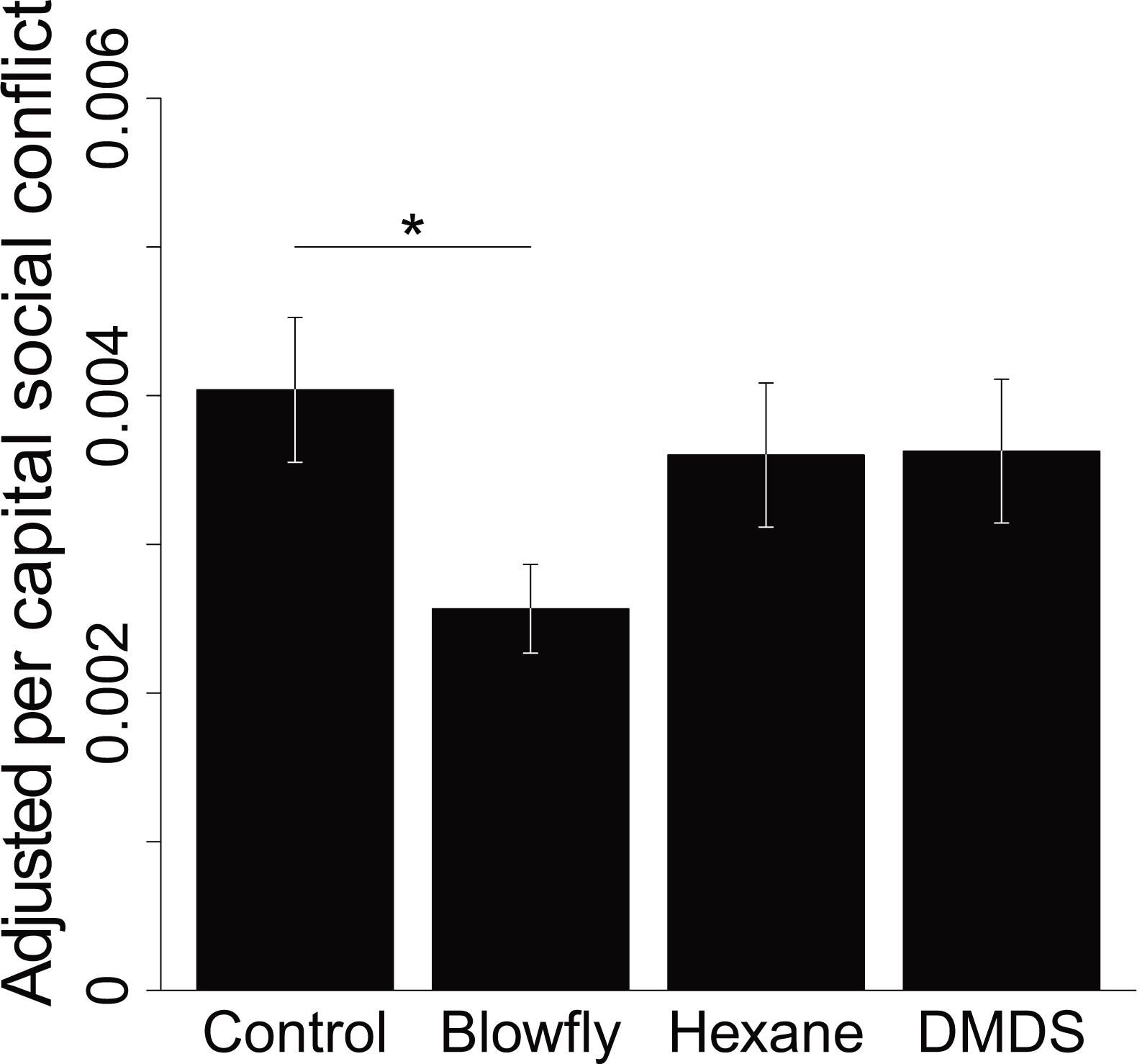
Adjusted per capita social conflict in different experimental treatments. Adjusted per capita social conflict (i.e. per capita social conflict divided by the total cooperative investment time) was lower in the blowfly than control treatments. There was no difference in adjusted per capita social conflict between the DMDS and hexane control treatments. * *P* < 0.05.

